# A *Wars2* mutant mouse shows a sex and diet specific change in fat distribution, reduced food intake and depot-specific upregulation of WAT browning

**DOI:** 10.1101/2022.05.23.493147

**Authors:** Milan Mušo, Liz Bentley, Lucie Vizor, Marianne Yon, Keith Burling, Peter Barker, Louisa A K Zolkiewski, Roger D Cox, Rebecca Dumbell

## Abstract

1.

**Background:** Increased waist-to-hip ratio (WHR) is associated with increased mortality and risk of type 2 diabetes and cardiovascular disease. The *TBX15*-*WARS2* locus has consistently been associated with increased WHR. Previous study of the hypomorphic *Wars2*^*V117L/V117L*^ mouse model found phenotypes including severely reduced fat mass, and white adipose tissue (WAT) browning, suggesting *Wars2* could be a potential modulator of fat distribution and WAT browning.

**Methods:** To test for differences in browning induction across different adipose depots of *Wars2*^*V117L/V117L*^ mice, we measured multiple browning markers of a 4-month old chow-fed cohort in subcutaneous and visceral WAT and brown adipose tissue (BAT). To explain previously observed fat mass loss, we also tested for the upregulation of plasma mitokines FGF21 and GDF15 and for differences in food intake in the same cohort. Finally, to test for diet-associated differences in fat distribution, we placed *Wars2*^*V117L/V117L*^ mice on low-fat or high-fat diet (LFD, HFD) and assessed their body composition by Echo-MRI and compared terminal adipose depot weights at 6 months of age.

**Results:** The chow-fed *Wars2*^*V117L/V117L*^ mice showed more changes in WAT browning marker gene expression in the subcutaneous inguinal WAT depot (iWAT) than in the visceral gonadal WAT depot (gWAT). These mice also demonstrated reduced food intake and elevated plasma FGF21 and GDF15, and mRNA from heart and BAT. When exposed to HFD, the *Wars2*^*V117L/V117L*^ mice showed resistance to diet-induced obesity and a male and HFD-specific reduction of gWAT : iWAT ratio.

**Conclusion:** Severe reduction of *Wars2* gene function causes a systemic phenotype which leads to upregulation of FGF21 and GDF15, resulting in reduced food intake and depot-specific changes in browning and fat mass.

## 2. Introduction

Increased waist-to-hip ratio (WHR) is associated with increased mortality and risk of coronary heart disease, myocardial infarction and type 2 diabetes (Vazquez *et al*., 2007; Snijder *et al*., 2003; Wang *et al*., 2005; Canoy, 2008; Mason, Craig and Katzmarzyk, 2008; Myint *et al*., 2014; Emdin *et al*., 2017; Peters, Bots and Woodward, 2018). The most recent meta-analysis identified 346 different loci associated with WHR adjusted for body mass index (WHRadjBMI) with most of the candidate genes being enriched in adipocytes and multiple fat depots (Pulit *et al*., 2018).

The *TBX15-WARS2* locus, which spans ∼1Mb and includes genes *TBX15, WARS2* and regions downstream of *SPAG17*, is consistently associated with WHR across multiple meta-analyses (Heid *et al*., 2010; Shungin *et al*., 2015; Pulit *et al*., 2018). Since the majority of SNPs in this region overlap the non-coding part of the genome, the effector genes remain to be identified (Maurano *et al*., 2012; Mušo *et al*., 2020). WARS2 is a mitochondrial tryptophanyl-tRNA synthetase, recently associated with angiogenesis and brown adipose tissue metabolism (Wang *et al*., 2016; Pravenec *et al*., 2017). Expression of both *TBX15* and *WARS2* in subcutaneous adipose tissue was associated with multiple metabolic traits including BMI and Matsuda insulin sensitivity index (Civelek *et al*., 2017). The GTEx database links the *TBX15-WARS2* locus risk SNPs to the expression of *WARS2* in multiple human tissues, but a few studies have also linked the locus to *TBX15* expression in adipose (Heid *et al*., 2010; GTEx-Consortium, 2013; Civelek *et al*., 2017).

Our group has previously established a *Wars2*^*V117L/V117L*^ mouse model where a N-ethyl-N-nitrosourea (ENU)-induced hypomorphic mutation causes defective splicing and results in only 0-30% of the full-length protein remaining across different tissues (Agnew *et al*., 2018). Homozygous *Wars2*^*V117L/V117L*^ mice showed mitochondrial electron transport chain (ETC) complex deficiency in multiple tissues, hypertrophic cardiomyopathy, sensorineural hearing loss and failure to gain fat mass. Importantly, white adipose tissue (WAT) showed upregulation of mitochondria and browning markers such as uncoupling protein 1 (UCP1) and mRNA levels of cell death-inducing DNA fragmentation factor subunit alpha (DFFA)-like effector a (*Cidea)* and iodothyronine deiodinase 2 (*Dio2*) genes. On the other hand, the brown adipose tissue (BAT) was dysfunctional and showed reduced browning marker expression. Elevated serum fibroblast growth factor-21 (FGF21) and mRNA from heart, muscle and white adipose suggested a mechanism by which at least part of the browning in adipose tissue may be mediated systemically (Fisher *et al*., 2012).

Another mitokine frequently co-induced with FGF21 in response to mitochondrial stress is growth/differentiation factor 15 (GDF15). GDF15 was previously reported to be an inducer of taste aversion and a suppressor of food intake by acting in the hindbrain where its receptor GDNF family receptor α–like (GFRAL) is expressed (Patel *et al*., 2019, Mullican *et al*., 2017). We hypothesised that a possible elevation of GDF15 levels could be thus affecting food intake and in effect the fat mass in *Wars2*^*V117L/V117L*^ mice.

In this follow-up study, we set out to explore whether WARS2 could be a regulator of white adipose browning and fat distribution. We initially tested whether the previously observed WAT browning effects in *Wars2*^*V117L/V117L*^ mice differed between different depots and whether changes in FGF21, GDF15 and food intake are observed and thus could explain the failure to gain fat mass in the chow-fed mice. Given that human polymorphisms in the *TBX15-WARS2* locus are associated with a less severe reduction in *WARS2* expression (GTEx-Consortium, 2013), we included heterozygous *Wars2*^*+/V117L*^ mice in our study. We evaluated the effect of high- and low-fat diet challenge (HFD – 60% kcal fat, LFD – 10% kcal fat) on adiposity and tested for any diet and depot specific differences in fat mass loss.

## 3. Materials and methods

### Animal models

All mice used in this study were housed in the Mary Lyon Centre at MRC Harwell. Mice were kept and studied in accordance with UK Home Office legislation and local ethical guidelines issued by the Medical Research Council (Responsibility in the Use of Animals for Medical Research, July 1993; Home Office license 30/3146 and 30/3070). Procedures were approved by the MRC Harwell Animal Welfare and Ethical Review Board (AWERB). Mice were kept under controlled light (light 7am–7pm, dark 7pm–7am), temperature (21 ± 2°C) and humidity (55 ± 10%) conditions. They had free access to water (9–13 ppm chlorine) and were fed *ad libitum* on a commercial chow diet (SDS Rat and Mouse No. 3 Breeding diet, RM3, 3.6 kcal/g) unless stated otherwise. Mice were group housed unless stated otherwise and were randomised into sex-matched cages on weaning. Researchers were blinded to the genotype of mice until analysis of the data.

### Experiment 1 - Molecular and hormonal investigation of *Wars2*^*V117L/V117L*^ mice

*Wars2*^*V117L/V117L*^ mice were generated and genotyped as previously described (Potter *et al*., 2016; Agnew *et al*., 2018). Tissues and plasma were collected in experiments previously described (Agnew *et al*., 2018). Briefly, 4-month-old male and female *Wars2*^*V117L/V117L*^and *Wars2*^*+/+*^ mice (n = 5-7) were humanely killed by terminal anaesthesia, and retro-orbital blood was collected into lithium-heparin microvette tubes (CB300, Sarstedt, Numbrecht, Germany). Death was confirmed by cervical dislocation and mice were then dissected and kidney, liver, muscle, heart, iWAT, gWAT and BAT collected. Tissues were directly placed in cryotubes and snap frozen in liquid nitrogen and samples were stored at −70°C before subsequent analyses by qPCR.

### Experiment 2 - Food intake measurements in *Wars2*^*V117L/V117L*^ mice

Four-week-old male and female *Wars2*^*V117L/V117L*^, *Wars2*^*+/V117L*^, and *Wars2*^*+/+*^ mice were pair-housed by genotype with *ad libitum* access to RM3 diet (n = 4 – 10 cages). Food was weighed twice a week until 16 weeks of age, and mice were weighed weekly. The mean weekly food intake per week per cage was calculated and cumulative food intake analysed.

### Experiment 3 - Body fat distribution in *Wars2*^*V117L/V117L*^ mice on HFD

We investigated body composition and fat distribution in male and female *Wars2*^*V117L/V117L*^, *Wars2*^*+/V117L*^, and *Wars2*^*+/+*^ mice challenged with a high fat diet (HFD). Experimental cohort numbers were based on estimates made using GraphPad Statmate using gWAT : iWAT ratios from previous experiments. We generated three cohorts of males and females, which were weaned directly onto HFD (*Research Diets*, D12450J) or matched low-fat diet (LFD, *Research Diets*, D12492) (n = 9 – 22, 185 mice in total).

Total body mass was measured every two weeks from 4 weeks of age on a scale calibrated to 0.01g. Body composition of the mice was measured every two weeks using an Echo-MRI (EMR-136-M, Echo-MRI, Texas, USA). The readings were total fat mass (g) and total lean mass (g). At 24 weeks old, mice were humanely killed by cervical dislocation and tissues were dissected and individual fat depots were dissected and weighed: interscapular BAT (iBAT), interscapular WAT (isWAT), perirenal BAT (prBAT), perirenal WAT (prWAT), inguinal WAT (iWAT), gonadal WAT (gWAT), mesenteric WAT (mWAT), and epicardial WAT (cWAT). gWAT: iWAT ratio was calculated from these weights as an indicator of visceral : subcutaneous fat distribution as described in (Gray *et al*., 2006).

### Experiment 4 - Body weight and composition in heterozygous knockout *Wars2*^*+/-*^ mice on HFD

NIH KOMP *Wars2*^+/-^ mice (*Wars2*^*tm1(KOMP)Vlcg*^) obtained from the KOMP repository (https://www.komp.org/) were imported into our laboratory previously (Agnew *et al*., 2018). Female *Wars2*^*+/-*^ and *Wars2*^*+/+*^ mice were weaned directly onto HFD or LFD (n = 7-9, 32 mice) and maintained until 12 months when they were weighed, and body composition analysed by Echo-MRI (EMR-136-M, Echo-MRI, Texas, USA). At 12 months old, mice were humanely killed by cervical dislocation and tissues were dissected and individual fat depots were dissected and weighed as in experiment 3.

### Quantitative PCR

Total RNA from adipose tissues (experiment 1) was extracted using the Direct-zol™ RNA MiniPrep Plus kit protocol (Zymo research, #R2071). RNA was reverse-transcribed using the SuperScript™ III Reverse Transcriptase Kit (ThermoFisher) to generate 2 μg of cDNA. mRNA gene expression was assayed using the TaqMan system (ThermoFisher) with the TaqMan FAM dye-labeled probes (Applied Biosystems, Invitrogen, U.S.A.) according to manufacturer protocols. Assays were carried out using an ABIPRISM 7500 Fast Real-Time PCR System (Applied Biosystems) and quantitation by the comparative C_T_ (ΔΔC_T_) analysis. Data was normalised to a geometric mean of 2 house-keeping genes specific to each tissue.

A mouse GeNORM analysis (PrimerDesing) for 6-8 genes was used to determine the most stable house-keeping genes. Taqman probes used in this study: *Canx* (Mm00500330_m1), *Rpl13a* (Mm01612986_gH), *Wars2* (Mm04208965_m1), *Ywhaz* (Mm01722325_m1), *Cidea* (Mm0042554_m1), *Cox7a1* (Mm00438297_g1), *Dio2* (Mm00515664_m1), *Ucp1* (Mm01244861_m1), *Fgf21* (Mm00840165_g1), *β-klotho* (Mm00502002_m1), *Pgc1α* (Mm01208835_m1), *Pparα* (Mm00440939_m1), *Pparγ* (Mm01184322_m1), *Prdm16* (Mm00712556_m1).

### Mitochondrial DNA copy number assay

Mitochondrial content in adipose tissue was assessed by ratio of mitochondrial DNA (mtDNA) to genomic DNA (gDNA) as assessed using qRT-PCR. Total DNA, which contains both gDNA and mtDNA, was extracted from adipose tissue (experiment 1) using the Dneasy Blood and Tissue Kit (Qiagen, # 69504). We amplified both the mouse genomic gene *Glyceraldehyde 3-phosphate dehydrogenase* (*Gapdh)* and mouse mitochondrial gene *Mitochondrially encoded NADH:Ubiquinone oxidoreductase core subunit 1 (mt-Nd1)* as proxies for genomic and mitochondrial DNA, respectively. Quantitative PCR was performed with 10ng DNA per reaction and 5 μM of each primer, using the Fast SYBR Green System on a ABIPRISM 7500 Fast Real-Time PCR Machine (Applied Biosystems). All samples were run in technical triplicates. Primers: *mt-Nd1*-Fw (CCCATTCGCGTTATTCTT), *mt-Nd1*-Rv (AAGTTGATCGTAACGGAAGC), *Gapdh-Fw (*CAAGGAGTAAGAAACCCTGGACC), *Gapdh*-Rv (CGAGTTGGGATAGGGCCTCT).

### Biochemical assays

Plasma fibroblast growth factor-21 (FGF21) levels were measured in blood plasma using Quantikine ELISA Mouse/Rat FGF21 Immunoassay (Quantikine, # MF2100). Mouse growth/differentiation factor 15 (GDF15) was measured using an in-house microtitre plate-based two-site electrochemiluminescence immunoassay using the MesoScale Discovery assay platform (MSD, Rockville, Maryland, USA). GDF-15 antibodies and standards were from R&D Systems (DuoSet # DY6385 BioTechne: Abingdon, UK).

### Statistical analysis

All statistical analyses were performed in Graph Pad Prism 9. Data outliers were identified using ROUT and omitted as indicated in each figure legend. Normality of distribution was evaluated using D’Agostino & Pearson normality test. Data was transformed where necessary in order to normalise their distribution prior to statistical analysis and details of the statistical tests used are described in each figure legend. Area under the curve for bodyweight, fat and lean mass was calculated in PRISM with Y=0 as a baseline. qPCR data was log-transformed and is shown as mean ± SD for visualisation and statistical analysis.

## 4. Results

### Browning is increased in both subcutaneous and visceral WAT depots of *Wars2*^*V117L/V117L*^ mice on chow diet

We set out to test whether browning effects previously observed in subcutaneous iWAT can also be observed in visceral gWAT, assessed by mRNA expression of a panel of browning and mitochondrial biogenesis gene markers in these mice at 4-months of age. In male iWAT of Wars2^*V117L/V117L*^ mice, as expected, we found increased expression of browning genes: *Cidea* increased by 0.61 ± 0.25 logFC (P = 0.0343), cytochrome c oxidase polypeptide 7A (*Cox7a*) by 0.60 ± 0.25 logFC (P = 0.0414) and *Dio2* by 0.93 ± 0.30 logFC (P = 0.0133) in (**Fig. 1**A). Male gWAT showed 0.56 ± 0.13 logFC (P = 0.0025) and 0.53 ± 0.10 logFC (P = 0.0006) increase in mRNA levels of both *Cidea* and the master regulator of mitochondrial biogenesis peroxisome proliferator-activated receptor gamma coactivator 1-α (*Pgc1α*) in Wars2^*V117L/V117L*^ mice (**Fig. 1**B). In female iWAT of Wars2^*V117L/V117L*^ mice, *Cidea, Dio2, Pgc1α* and *Pparα* were increased by 0.49 ± 0.15, 0.47 ± 0.15, 0.50 ± 0.12 and 0.33 ± 0.11 logFC, respectively (P = 0.0122, 0.0130, 0.0026, 0.0179, respectively) (**Supp. Fig. 1A**). The expression of browning genes in female gWAT was highly variable and *Pgc1α* was the only significantly upregulated gene 0.51 ± 0.17 logFC (P = 0.0176) in Wars2^*V117L/V117L*^ mice (**Supp. Fig. 1B**). In agreement with previous findings, BAT showed the reverse effect with reduced mitochondrial DNA content in both sexes (**Supp. Fig. 2A**) and reduced expression of browning marker gene expression in both sexes in *Wars2*^*V117L/V117L*^ mice (**Supp. Fig. 2B-C**). We next assessed mitochondrial mass as another marker of browning. Using a qPCR assay targeting both mtDNA and gDNA genes, we observed a significant increase of 0.43 ± 0.08 logFC and 0.23 ± 0.08 logFC in mtDNA : gDNA in male *Wars2*^*V117L/V117L*^ iWAT (P = 0.0002) and gWAT (P = 0.0264), respectively in *Wars2*^*V117L/V117L*^ mice (**Fig. 1C**). No genotype driven difference was seen in female mice (**Supp. Fig. 1C**). Together this is evidence of increased WAT browning, observed on multiple levels (mRNA, protein, mtDNA) in both iWAT and gWAT depots in *Wars2*^*V117L/V117L*^ mice with the specific effects differing between sexes and iWAT generally showing higher differences in fold change.

**Fig. 1.**
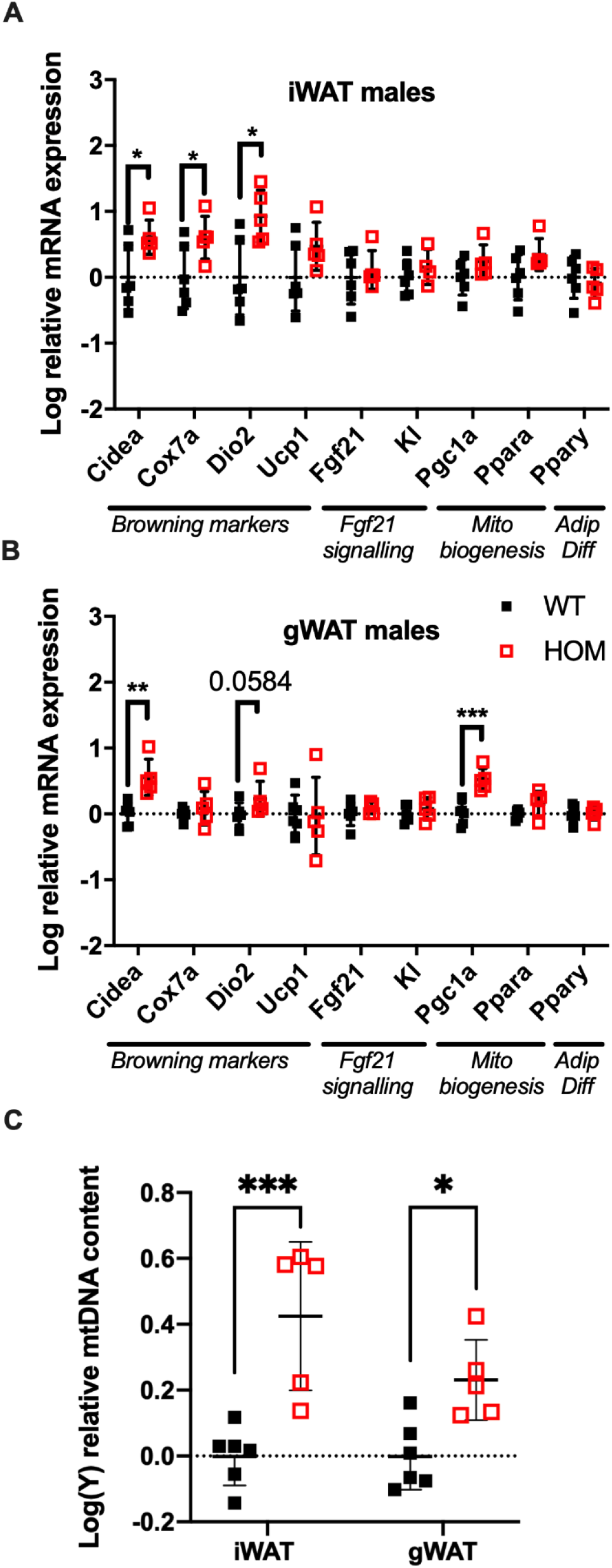

### Mitokines FGF21 and GDF15 are elevated in the *Wars2*^*V117L/V117L*^ mice on chow diet

12-month old *Wars2*^*V117L/V117L*^ mice were previously shown to have mitochondrial ETC complex deficiencies in multiple tissues and elevated plasma levels of the mitokine, FGF21 which may at least partially explain the WAT browning. We thus decided to measure circulating FGF21 and the appetite-suppressing mitokine GDF15 in free fed 4-month old mice (Patel *et al*., 2019). We observed an overall genotype effect (P = 0.0485) on FGF21 levels, with a 86% increase (P = 0.0364) in female *Wars2*^*V117L/V117L*^ mice and a non-significant trend for increase in males (**Fig. 2A**). GDF15 was significantly increased in *Wars2*^*V117L/V117L*^ mice of both sexes, with a 112% increase (P = 0.0014) in males and 158% increase (P = 0.0001) in females (**Fig. 2B**). We followed with a qPCR study of multiple tissues to show that *Fgf21* expression was elevated by 1.80 ± 0.13 mean difference of log10-fold change (logFC) ± (SE) (P = 0.0012), 0.38 ± 0.11 logFC (P = 0.0055), 0.47 ± 0.16 logFC (P = 0.0157), 0.41 ± 0.18 logFC (P = 0.0447) in the heart, BAT, muscle and kidney of *Wars2*^*V117L/V117L*^ mice respectively (**Fig. 2C**). *Gdf15* was elevated by 0.66 ± 0.09 of logFC (P<0.0001) and 0.84 ± 0.12 logFC (P<0.0001) in the heart and BAT, respectively (**Fig. 2D**). We also tested for changes in *Atf4* levels, one of the upstream regulators of *Gdf15* and *Fgf21*, but found no difference in any of the tissues (**Supp. Fig. 3**).

**Fig. 2.**
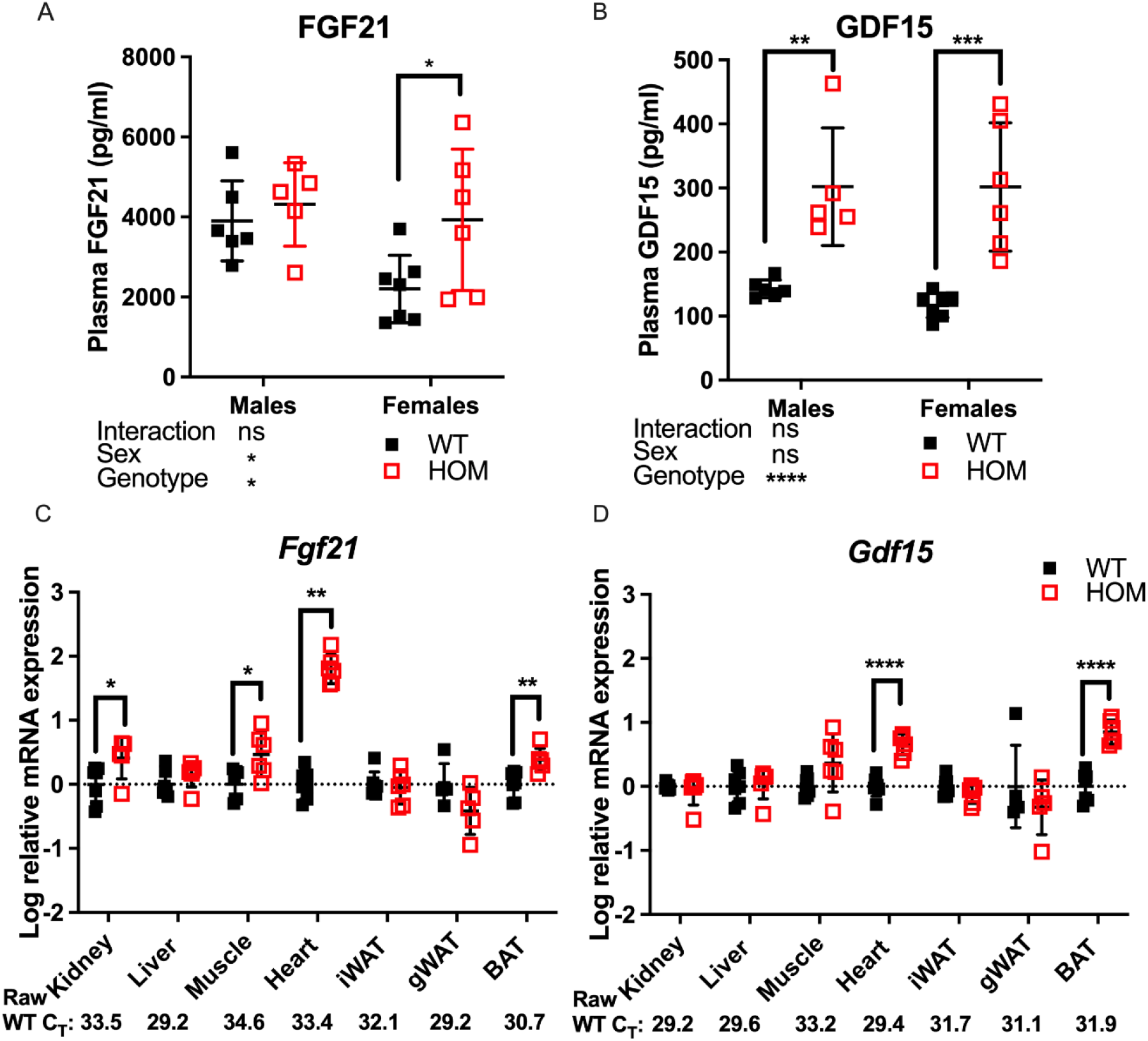

### Food intake is reduced in *Wars2*^*V117L/V117L*^ mice on a chow diet

We hypothesised that the elevated GDF15 levels may be contributing to reduced food intake in *Wars2*^*V117L/V117L*^ mice. To test an effect on food intake, we set up an independent cohort of pair-housed mice on regular RM3 chow diet. Male *Wars2*^*V117L/V117L*^ mice showed reduced cumulative food intake compared to wild-type mice already from the first timepoint at 7 weeks of age (P = 0.045) (**Fig. 3A-B**). Female *Wars2*^*V117L/V117L*^ mice showed significantly lower food intake from 10 weeks onwards (P = 0.0148). At 14 weeks of age, the male and female cumulative food intake was 17% (P = 0.0016) and 8.4% lower than wild-type (P = 0.0020), respectively (**Fig. 3A-B**). This is thus likely to have contributed to the lower bodyweight seen in these mice (**Fig. 3C-D**).

**Fig. 3.**
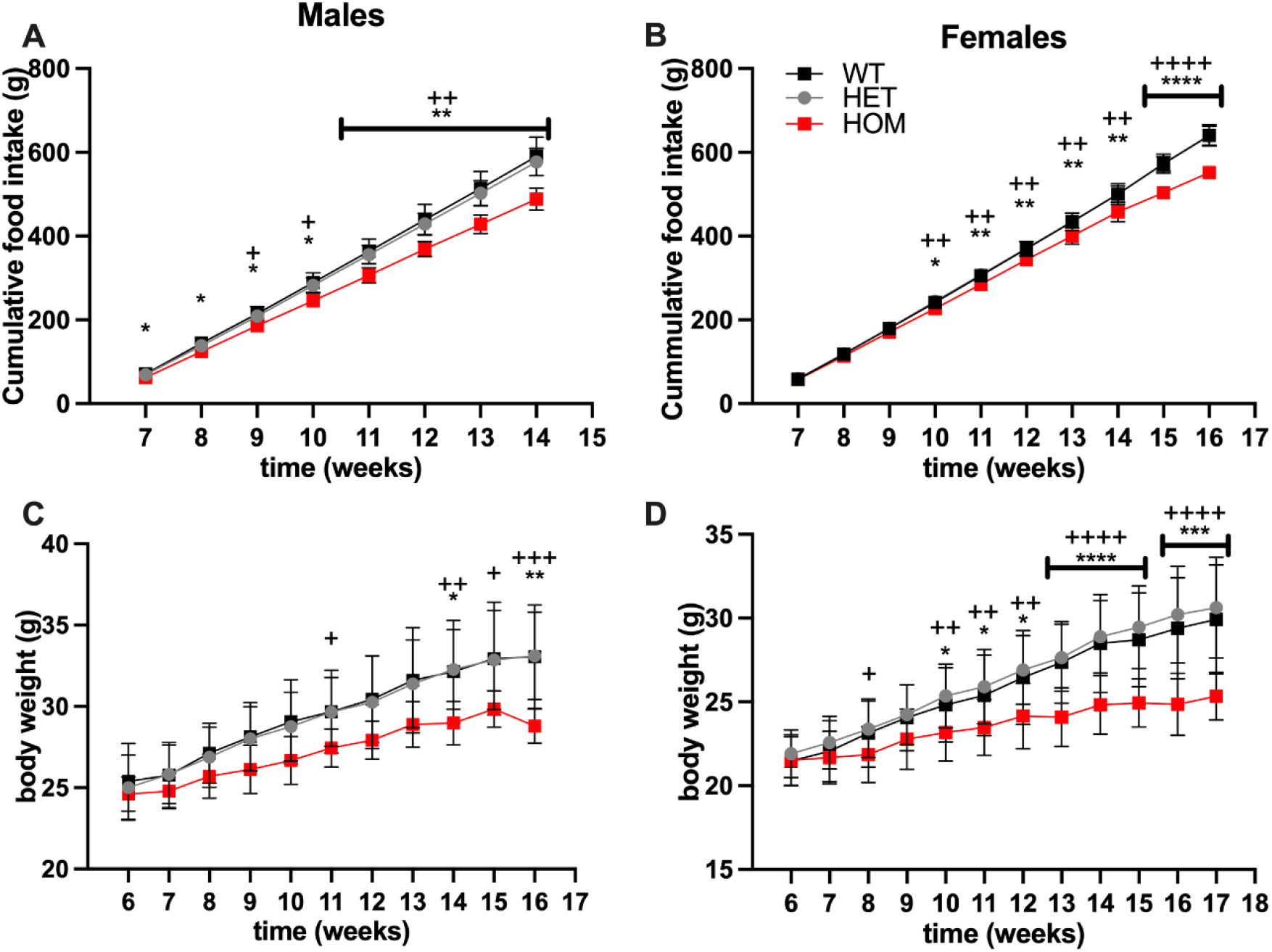

### Homozygous *Wars2*^*V117L/V117L*^ mice fail to gain fat mass due to growth and high-fat diet

In data from a small cohort of 6-month old Wars2^*V117L/V117L*^ males we previously showed a trend towards increased ratio of gWAT : iWAT mass (Agnew *et al*., 2018). To investigate whether an altered diet could reveal a fat distribution phenotype or whether it would alleviate the failure to gain fat mass found in these mice, *Wars2*^*V117L/V117L*^, *Wars2*^*+/V117L*^ and *Wars2*^*+/+*^ mice were placed on HFD and matched LFD. As expected, HFD increased body weight and fat mass in wild-type *Wars2*^*+/+*^ (week 24, males: P < 0.0001, P < 0.0001; females: P < 0.0001, P < 0.0001, respectively) and heterozygous *Wars2*^*+/V117L*^ mice (week 24, males: P = 0.0363, P < 0.0001, females: P < 0.0001, P < 0.0001, respectively). However, no significant effect of HFD on body weight was observed in *Wars2*^*V117L/V117L*^ mice of either sex (**Fig. 4A-B, Fig. 5A-B**). On LFD, significant bodyweight differences between wild-type and *Wars2*^*V117L/V117L*^ were observed and persisted from 14 (P = 0.0126) and 16 weeks of age (P = 0.0176) for males and females, respectively. On a HFD, significance was reached earlier, at 6 (P = 0.0016) and 12 (P<0.0001) weeks of age, respectively. Similar effects were observed between wildtype and homozygous mice for fat mass, which was significant from 6 and 12 weeks (male) and 10 and 16 weeks (female), six weeks earlier on HFD than on LFD respectively (**Fig. 4C-D, Fig. 5C-D**). Significant differences were also observed in the lean mass of *Wars2*^*V117L/V117L*^ mice, but these were of a smaller magnitude (**Fig. 4E-F, Fig. 5E-F**). When analysed over the time course using area under curve, these differences were maintained (**Supplementary Table 1**). In summary, most of the weight differences in *Wars2*^*V117L/V117L*^ mice were due to the reduction in fat mass and administering a high-fat diet exacerbated these differences.

**Fig. 4.**
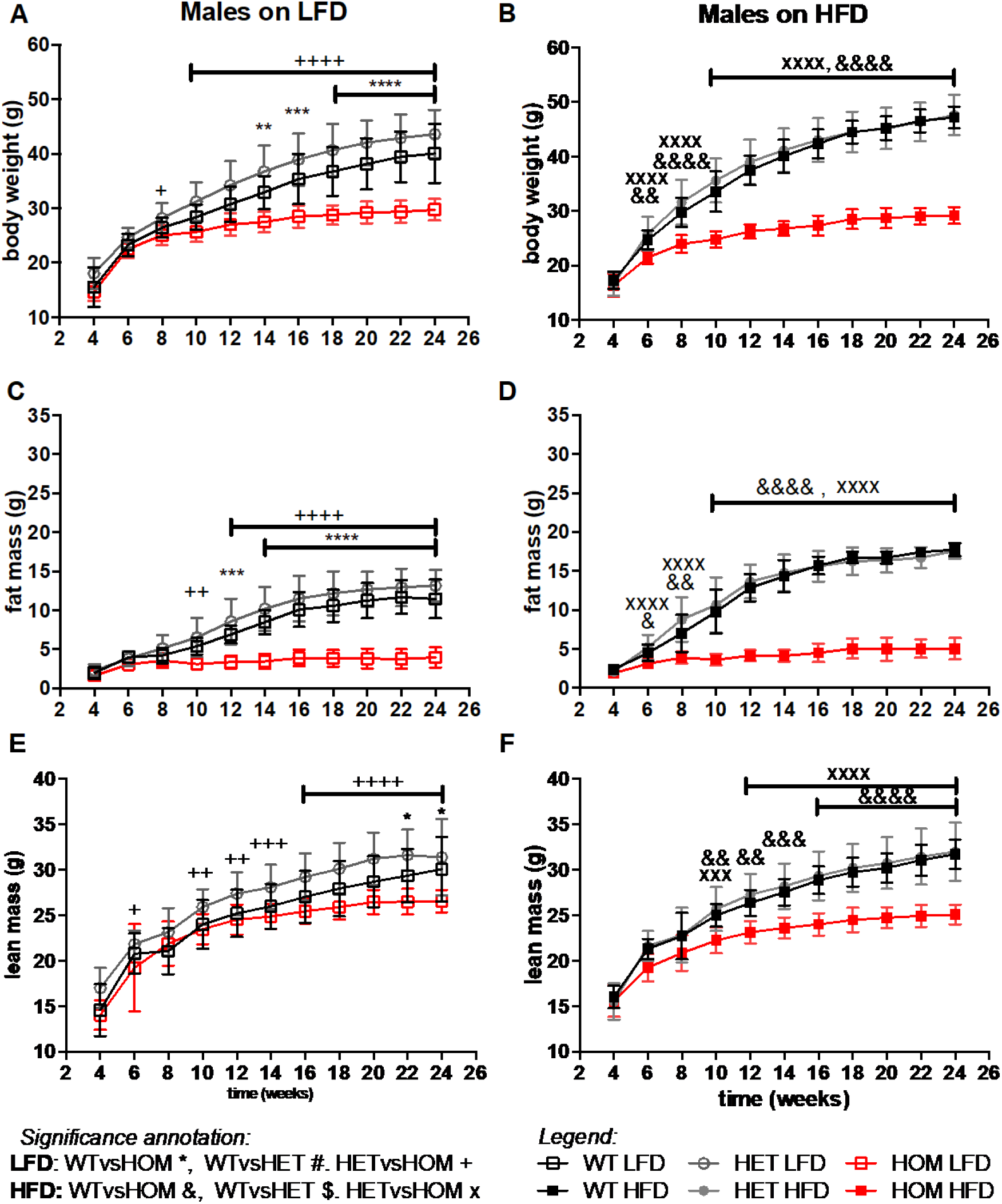

**Fig. 5.**
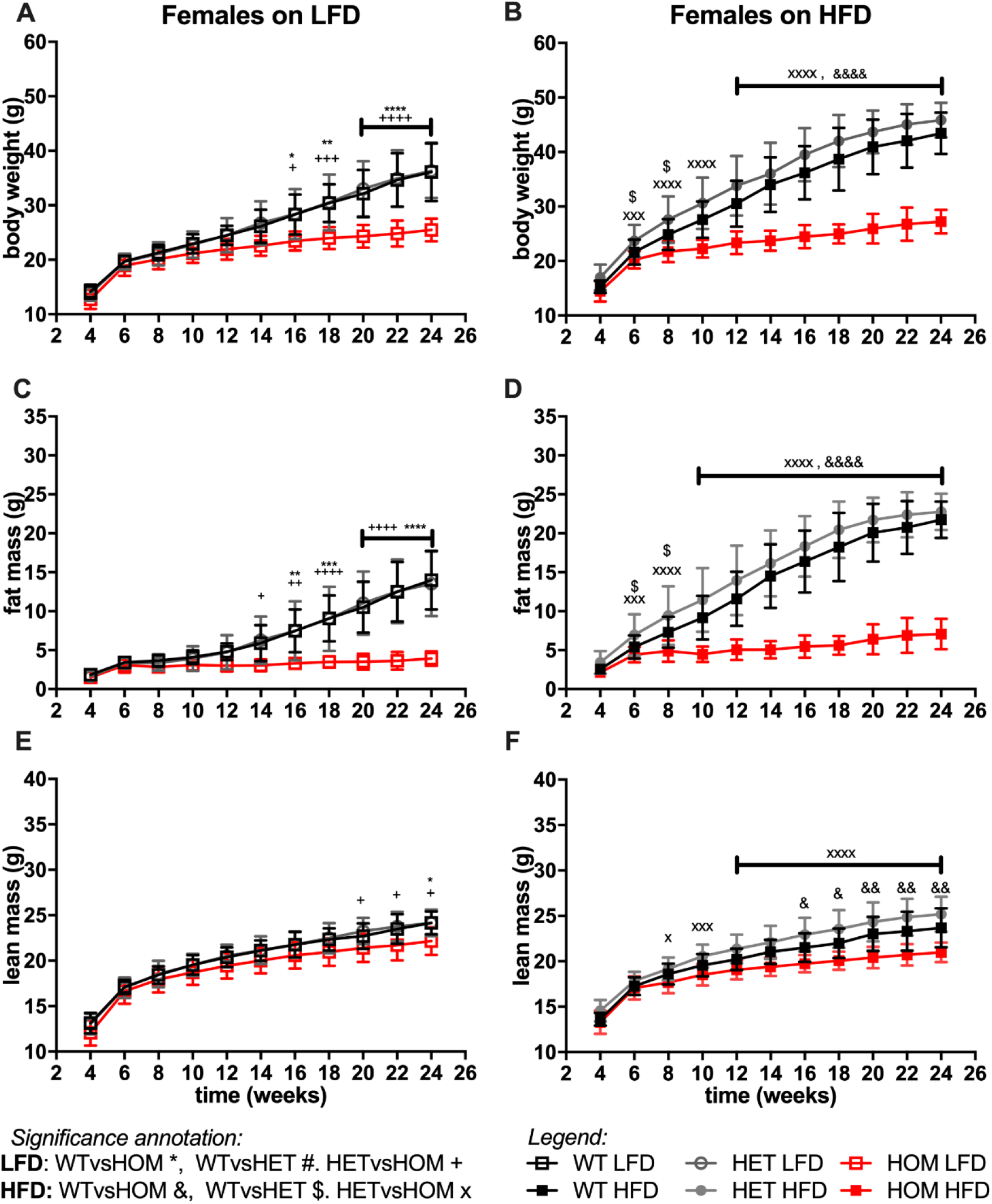

For all 3 measures, heterozygous *Wars2*^*+/V117L*^ mice also showed significant differences to *Wars2*^*V117L/V117L*^ mice at an earlier age than for wild type mice (**Fig. 4A-F, Fig. 5A-F**). A significant increase in bodyweight (P = 0.0476, P = 0.0416) and fat mass (P = 0.0418, P = 0.0430) of *Wars2*^*+/V117L*^ females on HFD was observed compared to wild-type mice at 6 and 8 weeks of age respectively, but this change did not persist in later timepoints. In line with this, 12-month-old heterozygous female knockout *Wars2*^*+/-*^ mice did not show any differences in body weight or composition on either diet (**Supp. Fig. 6**). In summary, we did not observe any reproducible differences between the heterozygous *Wars2*^*+/V117L*^ or *Wars2*^*+/-*^ mice and the wild-type mice.

### *Wars2*^*V117L/V117L*^ mice show reduction in the weights of multiple fat depots and a HFD and male specific elevation in gWAT : iWAT ratio

Since the majority of the weight differences in the *Wars2*^*V117L/V117L*^ mice could be explained by fat mass, we next evaluated differences in fat distribution by weighing fat depots from 24 week old mice, and we considered the ratio of gWAT : iWAT mass, (**Fig. 6, Supp. Fig. 4, Supp. Fig. 5**). Indeed, almost all fat depots weighed less in *Wars2*^*V117L/V117L*^ compared to wild-type or heterozygous mice. The only exceptions were male HFD gWAT, female HFD perirenal BAT and female LFD perirenal WAT which did not differ from *Wars2*^*+/+*^ or *Wars2*^*+/V117L*^. The lack of weight change in male *Wars2*^*V117L/V117L*^ gWAT on HFD together with the 1.372 ± 0.1755g lower iWAT weight (P<0.0001) resulted in an increased gWAT : iWAT ratio (P<0.0001), indicating higher visceral to subcutaneous fat ratio (**Fig. 6A,C,E**) Interestingly, no such trend was replicated in females where both iWAT and gWAT depot weights were reduced, by 1.508 ± 0.2398g (P<0.001) and 1.684 ± 0.2521g (P<0.001), respectively (**Fig. 6B,D,F**). No significant differences were observed between the heterozygous and wild-type mice for any of the fat depots apart for male iWAT on a LFD (P<0.05). Similarly, fat depots of 12-month-old female heterozygous *Wars2*^*+/-*^ mice in a separate cohort, did not show any significant differences (**Supp. Fig. 7**). This demonstrates that *Wars2*^*V117L/V117L*^ mice have much lower fat mass which is unequally shared by different fat depots and results in male and HFD-specific increase in gWAT : iWAT ratio.

**Fig. 6.**
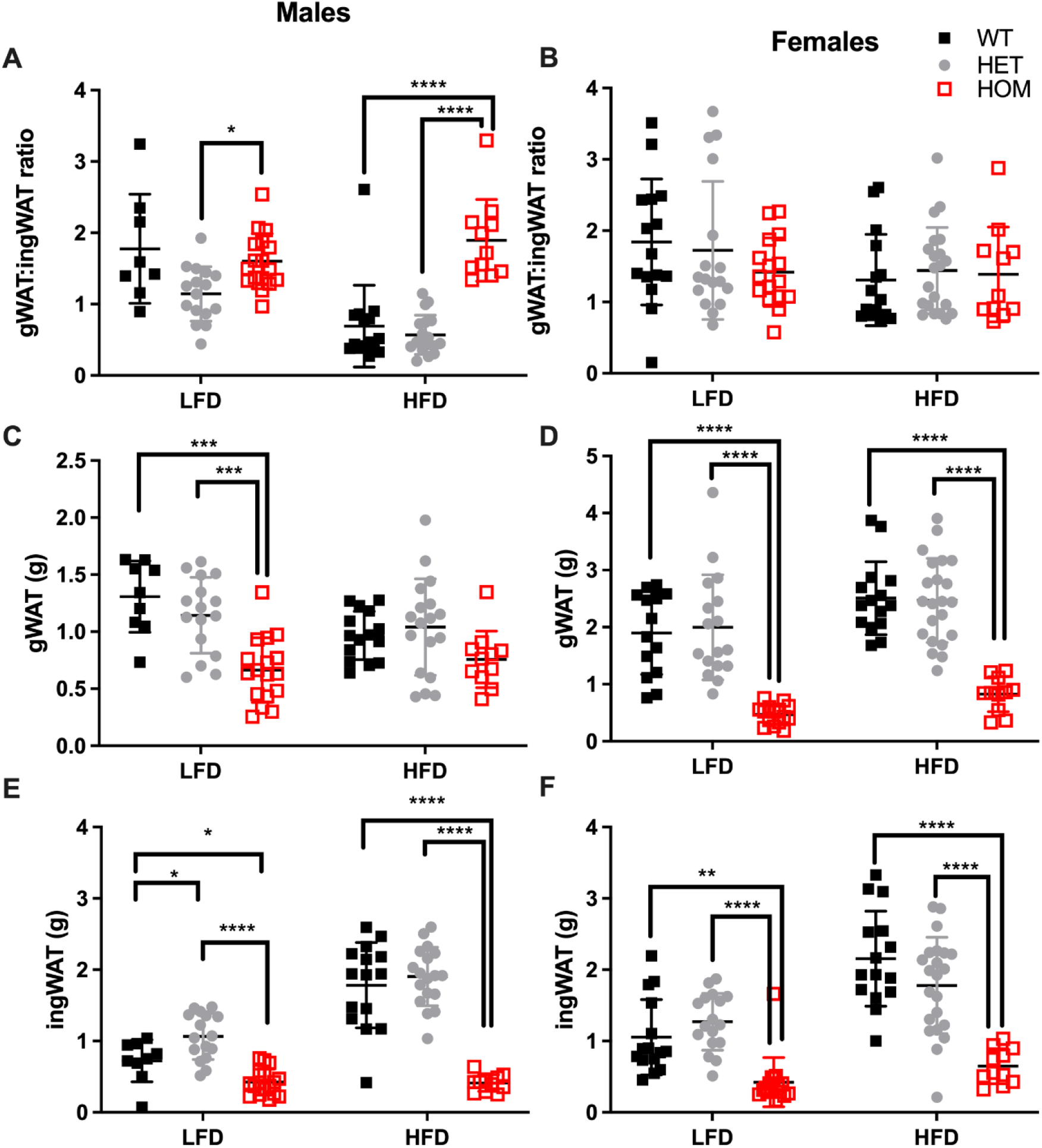

## 5. Discussion

We assessed fat depot differences in browning and showed that the magnitude of browning effects is greater in iWAT than in gWAT of chow-fed 4-month-old *Wars2*^*V117L/V117L*^ mice. This agrees with previous research which showed that gWAT has low browning marker expression and a very low browning capacity compared to iWAT(de Jong *et al*., 2015; Zuriaga *et al*., 2017). Our findings suggest that the adipose phenotypes in *Wars2*^*V117L/V117L*^ mice are driven systemically, secondary to a severe mitochondrial dysfunction in the heart, BAT and muscle. Firstly, we confirmed the upregulation of FGF21, an established inducer of WAT browning (Fisher *et al*., 2012; Agnew *et al*., 2018). Secondly, we showed higher plasma GDF15 in these mice which may contribute to the observed lower food intake that thus contributed to the reduced bodyweight and fat mass, as shown in other models of mitochondrial disease (Chung *et al*., 2017).

We have shown that *Wars2*^*V117L/V117L*^ mice fail to gain fat mass also when challenged with a HFD, accompanied by a male and HFD-specific upregulation of gWAT : iWAT ratio. This was likely driven by the lower mass of iWAT and the relatively unchanged visceral gWAT on HFD. In general, all other male visceral depots showed a reduction of fat mass in male *Wars2*^*V117L/V117L*^ mice. It would be interesting to extend these observations using other methods, such as small animal X-ray computed tomography (CT) system, that could accurately verify the effect on overall fat distribution over time (Sasser *et al*., 2012). This male-specific effect is in line with sexual dimorphism which is an established feature of fat distribution (Pulit, Karaderi and Lindgren, 2017). In fact, the *TBX15-WARS2* locus also contains an independent male-specific WHRadjBMI-association signal (Shungin *et al*., 2015). Further study will be required to explain the diet specificity. However, HFD was previously shown to induce browning and it could thus potentiate the depot-specific differences observed in chow-fed animals and thus contribute to HFD-specific fat mass loss seen in WAT and not gWAT (García-Ruiz *et al*., 2015).

Is it possible that a similar mechanism relating mitochondrial failure in the heart and other tissues together with WAT browning might drive the WHR signal in humans? Indeed, rare variants in genes of the mitochondrial genomes and in another member of the *ARS2* family, *DARS2*, have all been associated with WHR (Justice *et al*., 2019). Furthermore, in the Common Metabolic Diseases Knowledge Portal, the *TBX15-WARS2* locus is associated with cardiovascular traits such as stroke severity and peripheral vascular disease in people with type 2 diabetes (cmdgenkp.org, no date a), whilst variants in the *WARS2* gene are linked to diastolic blood pressure (cmdgenkp.org, no date b). This suggests that a systemic mechanism could explain the WHR GWAS association in humans.

In conclusion, we have shown that a hypomorphic mutation in the *Wars2* gene causes a severe failure to gain body mass and results in changes to fat distribution in male mice on a HFD. We also reveal differences in browning propensity of different WAT depots and elevation of FGF21 and GDF15 which likely partly explain some of these phenotypes. These data support a potential functional role for *WARS2* in the WHRadjBMI *TBX15-WARS2* locus, which could be further investigated in human studies where *WARS2* expression varies by genotype.

## Author Contributions

MM, RDC and RD designed and supervised the experiments, analysed data, prepared figures, and wrote the manuscript with input from all authors. MM carried out the molecular studies and body composition measurements. Food intake analysis was carried out by LB, MM and LV. Cohorts were managed by LB and LV. Fat depot weight measurements were performed by MY, EF, HN, RD and MM. LZ assisted with molecular studies. KB and PB carried out the GDF15 ELISAs.

## Acknowledgements

We thank Emily File in the MLC for excellent technical assistance.

## Contribution to the Field

Increased waist-to-hip ratio is associated with increased mortality and risk of coronary heart disease, myocardial infarction and type 2 diabetes. Although multiple theories exist to explain these associations, it is not yet certain why altered fat distribution would increase disease risk and whether targeting some of its molecular pathways could be utilised for disease prevention. Large genome wide association studies have uncovered hundreds of genomic loci, but for most of these, the causal gene is largely unknown. In this study, we have tested one candidate gene *WARS2* in the *TBX15/WARS2* locus by studying a mouse model with a damaging mutation in this gene. This mouse model has been previously shown to fail to gain fat mass and showed a complex phenotype with cardiomyopathy and increased white adipose tissue browning. Here, for the first time we also show that these mice show a diet and sex specific difference in fat distribution and thus implicate *WARS2* as a potential modulator of fat distribution. Since coding mutations in other mitochondrial protein genes have been associated with waist-to-hip ratio, this finding strengthens the link between mitochondrial function and adipose biology.

## Figure captions

**Fig. 7.** Increased browning in inguinal WAT (iWAT) and gonadal WAT (gWAT) of 4-month old male *Wars2*^*V117L/V117L*^ mice. (A,B) Relative expression of browning, Fgf21 signalling, mitochondrial biogenesis and adipose differentiation markers in iWAT and gWAT, respectively. Normalised to geometric mean of *Canx* and *Ywhaz*. Data was log-transformed and assessed by unpaired t-test or Mann-Whitney test (iWAT for *Dio2* and *Fgf21*) based on their distribution, n = 6 and 5 wildtype and homozygotes respectively in iWAT and gWAT. (C) qPCR analysis of *mt-Nd1:Gapdh* ratio signifying mitochondrial : genomic DNA (mtDNA : gDNA) ratio. 2-way ANOVA with post-hoc comparison of genotypes, n = 5. All data shown as mean ± SD.

**Fig. 8.** GDF15 and FGF21 levels are elevated in 4-month old *Wars2*^*V117L/V117L*^ mouse plasma. ELISA analysis of FGF21 (A) and GDF15 (B) levels in males (n = 5-6) and females (n = 6-7). Analysis by 2-way ANOVA followed by post-hoc Sidak multiple comparison. qPCR analysis of *Fgf21* (C) and *Gdf15* (D) levels in multiple tissues from the female mice used in (A) and (B) (n = 5-7). Data was log-transformed and assessed by unpaired t-test or Mann-Whitney test (*Fgf21* in Heart). Mean raw C_T_ values are shown for wildtype tissues. All data shown as mean ± SD. Average WT C_T_ values listed beneath the graphs.

**Fig. 9.** Food Intake and bodyweight are reduced in *Wars2*^*V117L/V117L*^ mice. Cumulative food intake in (A) males (n = 4-10) and (B) females (n = 8-9). N represents 1 cage of 2 mice of the same genotype. Bodyweight in the same cohort of (C) males (n = 8-20) and (B) females (n = 12-18) where N represents each mouse. Significance at specific time points was calculated with 1-way ANOVA with multiple comparisons. Significance symbols for WT x HET:*, HET x HOM:+.

**Fig. 10.** *Wars2*^*V117L/V117L*^ mice fail to gain fat and lean mass during growth and due to high fat diet feeding. Three cohorts of 6-month old male (n = 9-18) mice on low-fat (LFD) or high-fat diet (HFD) were pooled and assessed for body weight (A and B), fat mass (C and D), and lean mass (E and F), respectively. Genotypes: *Wars2*^*+/+*^ (WT), *Wars2*^*+/V117L*^ (HET) and *Wars2*^*V117L/V117L*^ (HOM). For male mice one homozygote on a LFD and one wildtype on a HFD were excluded as outliers (identified using ROUT in GraphPad PRISM 9). Significance at specific time points was calculated with 2-way ANOVA with Tukey’s multiple comparison analysis for all groups. Significant difference between *Wars2*^*+/+*^ (WT) and *Wars2*^*V117L/V117L*^ (HOM) is shown as * p < 0.05, ** p< 0.01, *** p<0.001. Comparisons between other groups are depicted in the same way using the symbols (+, &, x, $, #) annotated in the top right corner.

**Fig. 11.** *Wars2*^*V117L/V117L*^ mice fail to gain fat and lean mass during growth and due to high fat diet feeding. Three-cohorts of 6-month old female (n = 11-22) mice on low-fat (LFD) or high-fat diet (HFD) were pooled and assessed for body weight (A and B), fat mass (C and D), and lean mass (E and F), respectively. Genotypes: *Wars2*^*+/+*^ (WT), *Wars2*^*+/V117L*^ (HET) and *Wars2*^*V117L/V117L*^ (HOM). Significance at specific time points was calculated with 2-way ANOVA with Tukey’s multiple comparison analysis for all groups within each sex. Significance between *Wars2*^*+/+*^ (WT) and *Wars2*^*V117L/V117L*^ (HOM) is shown as * p < 0.05, ** p< 0.01, *** p<0.001. Comparisons between other groups are depicted in the same way using the symbols (+, &, x, $, #) annotated in the top right corner.

**Fig. 12.** Gonadal to inguinal WAT (gWAT : iWAT) ratio is elevated in *Wars2*^*V117L/V117L*^ males on a HFD. gWAT : iWAT ratio was calculated for 6-month old male (n = 9-18) and female (n = 11-22) mice either on low fat (LFD) and high fat (HFD) diets (A, B). The individual gWAT (C, D) and iWAT (E, F) weights are shown below. To fit a normal distribution, male and female gWAT : iWAT ratio data and male iWAT data were transformed by Y=Log_2_(Y). The gWAT male and female data were normally distributed (D’Agostino & Pearson normality test) and the iWAT female data showed some deviation from normality (P = 0.0476). Significance was tested using 2-way ANOVA with Tukey’s multiple comparison test between genotype for each diet. Significant differences in multiple comparisons of WT, HET and HOM on each diet are depicted as *p < 0.05, **p< 0.01, ***p<0.001.

