## Supplementary Data for "A *Wars2* mutant mouse shows a sex and diet specific change in fat distribution, reduced food intake and depot-specific upregulation of WAT browning"

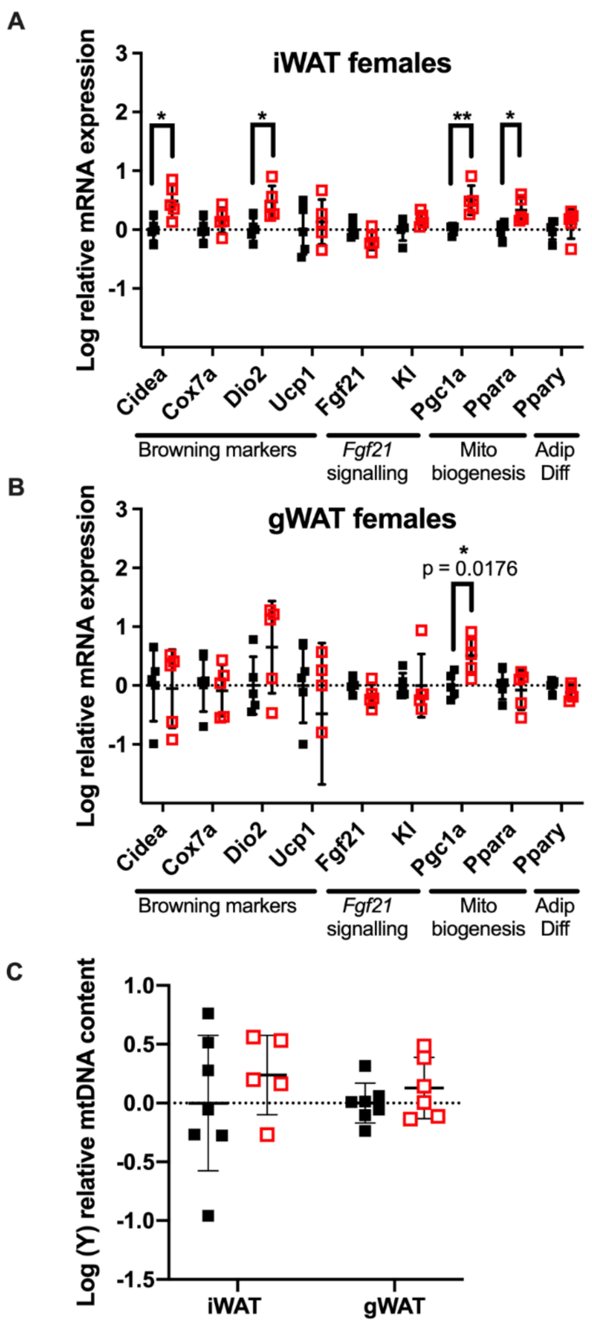

**Supp. Fig. 1 Increased browning in inguinal WAT (iWAT) and gonadal WAT (gWAT) of 4-month old female *Wars2*$^{V117L/V117L}$ mice.** (A,B) Relative expression of browning, Fgf21 signalling, mitochondrial biogenesis and adipose differentiation markers in iWAT and gWAT, respectively. Normalised to geometric mean of *Canx* and *Ywhaz*. Data was log-transformed and assessed by individual unpaired t-test's or a Mann-Whitney test (KI in iWAT, KI and UCP-1 in gWAT) based on the distribution (C) qPCR analysis of *mt-Nd1 : Gapdh* ratio signifying mitochondrial : genomic DNA (mtDNA : gDNA) ratio. 2-way ANOVA with Sidak's post-hoc comparison of genotypes, n = 5-6. All data shown as mean ± SD.

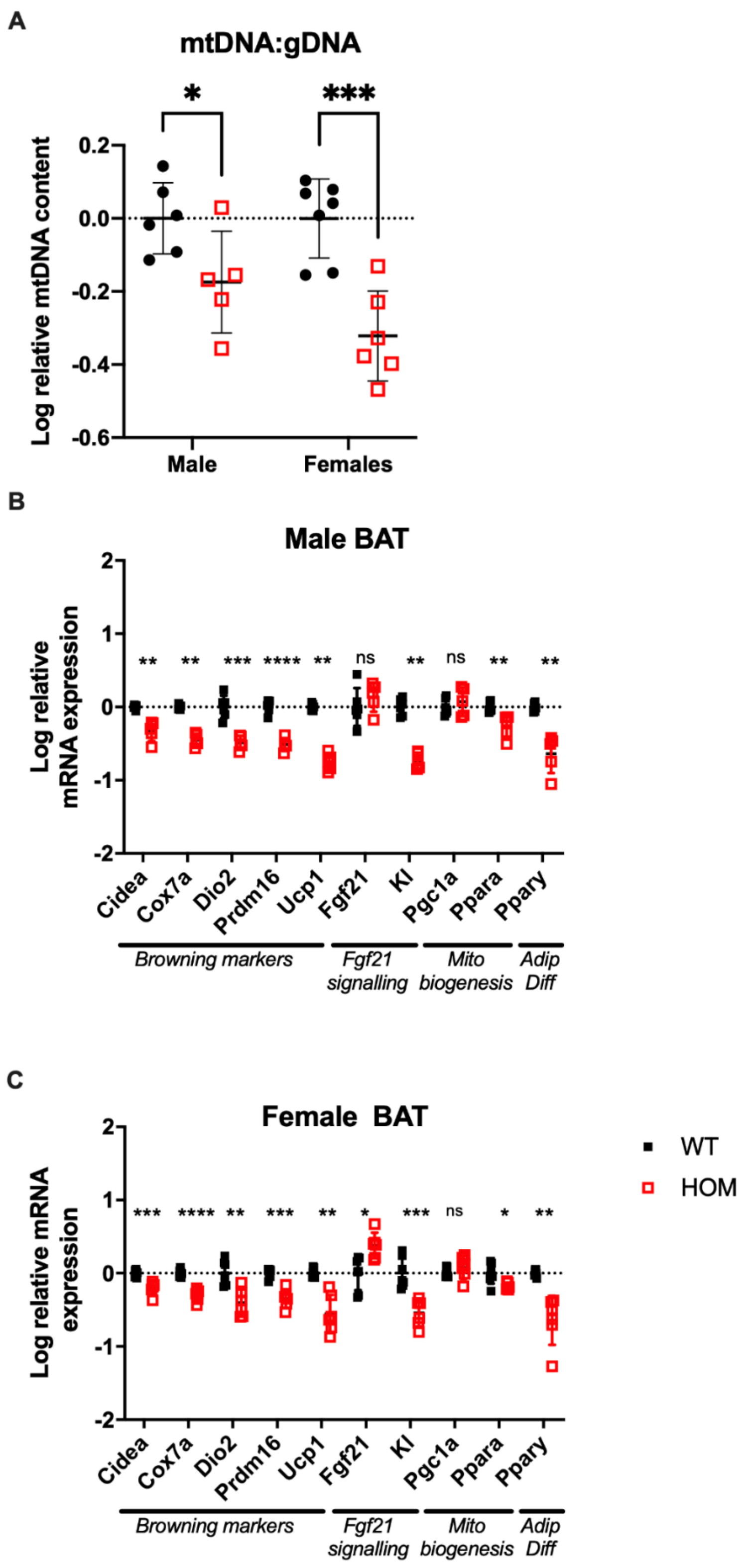

**Supp. Fig. 2** Reduced browning signature in interscapular brown adipose tissue (BAT) of 4-month old *Wars2*$^{V117L/V117L}$ mice. (A) qPCR analysis of *mt-Nd1:Gapdh* ratio signifying mitochondrial : genomic DNA (mtDNA :gDNA) ratio. Log(Y) transformed data was analysed by 2-way ANOVA with post-hoc (Sidak multiple comparison test) comparison of genotypes, n = 5-7. (B,C) Relative expression of browning, Fgf21 signalling, mitochondrial biogenesis and adipose differentiation markers in male (n = 6 wildtype and 5 homozygotes) and female BAT (n = 7 wildtypes and 6 homozygotes), respectively. Normalised to geometric mean of *Canx* and *Ywhaz*. Normality of distribution was evaluated using D'Agostino & Pearson normality test. Data was log-transformed and assessed by individual unpaired t-test (unless otherwise indicated), Welch's t-test (female *UCP-1*, *PPAR*$\alpha$ and *PPAR*$\gamma$, male *Cidea, PPAR*$\gamma$*,)* or Mann-Whitney test (male *Cox7a, UCP-1, KI*) depending on their distribution and variances. All data shown as mean ± SD.

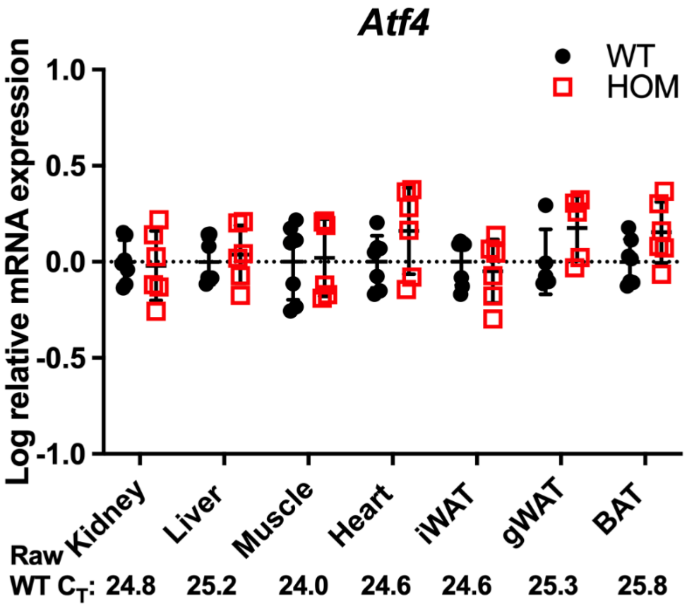

**Supp. Fig. 3** ***Atf4*** **expression is not altered in 4-month-old** ***Wars2***$^{V117L/V117L}$ **mouse plasma.** qPCR analysis in multiple tissues from female 4-month-old mice (n = 5-7). Data was log-transformed and assessed by unpaired t-test. All data shown as mean ± SD. Average WT $C_T$ values listed beneath the graphs.

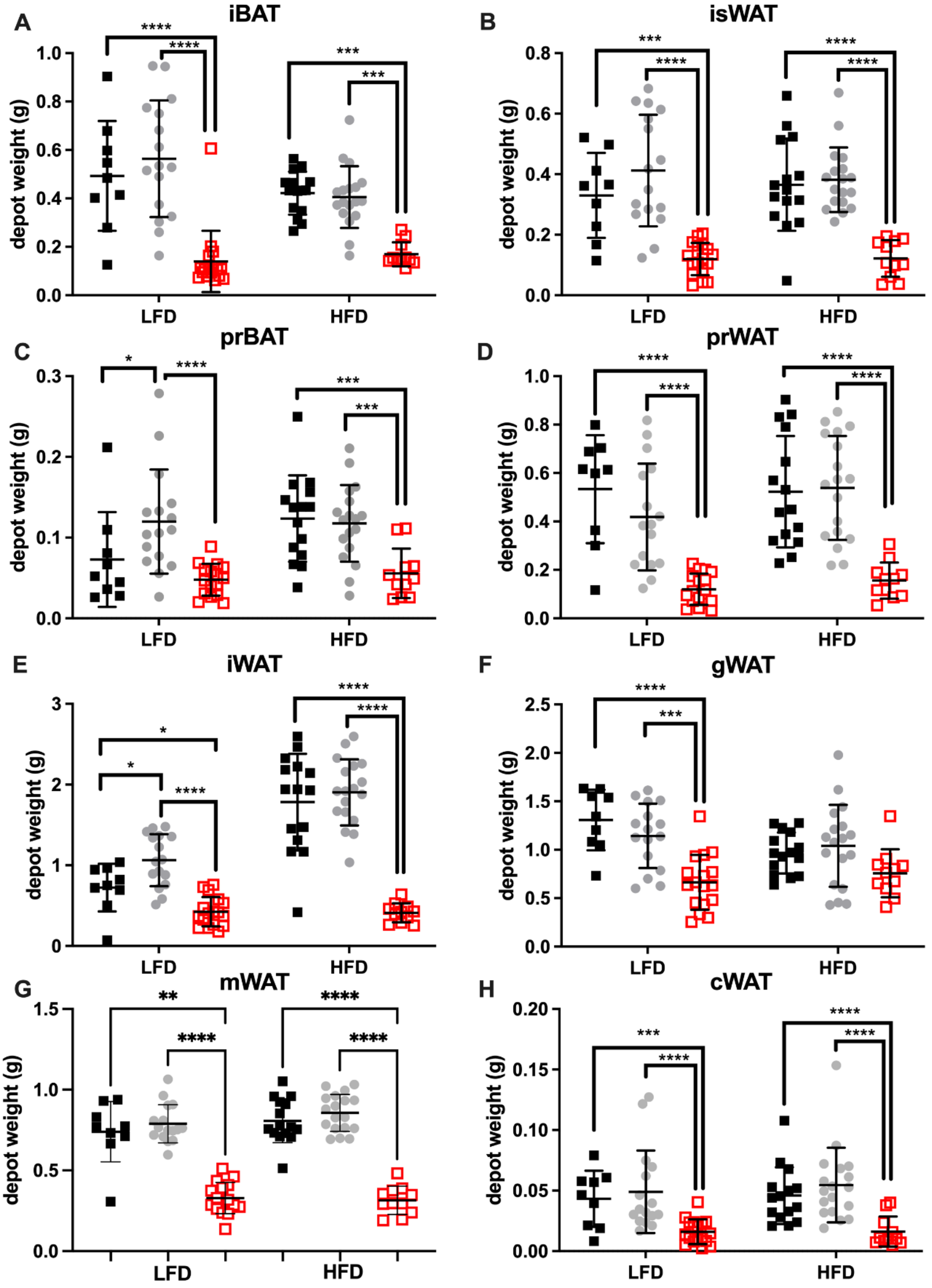

**Supp. Fig. 4 Multiple male *Wars2*$^{V117L/V117L}$ fat depots are affected.** Fat depots of 6-month old male (n = 9-18) mice on low fat (LFD) or high fat (HFD) diets were dissected and the following fat depots weighed: interscapular BAT (iBAT, A) , interscapular WAT (iWAT, B), perirenal BAT (prBAT, C), perirenal WAT (prWAT, D), inguinal WAT (iWAT, E gonadal

WAT (gWAT, F) mesenteric WAT (mWAT, G), cardiac WAT (cWAT, H). Data in E and F are from Figure 3 and are included here for comparison. Normality of distribution was evaluated using D'Agostino & Pearson normality test. In order to normalise distribution, data was Y=log2(Y) transformed for prBAT, prWAT, iWAT before analysis. Significance was tested using 2-way ANOVA with Tukeys multiple comparison test between all the groups (A-F, H). One outlier was identified and removed by ROUTE from the mWAT data set which was then analysed using a nonparametric Kruskal-Wallis Test and a Dunn's multiple comparisons test (G). Significant differences in multiple comparisons of WT, HET and HOM on each diet are depicted as $*p < 0.05$, $**p < 0.01$, $***p < 0.001$.

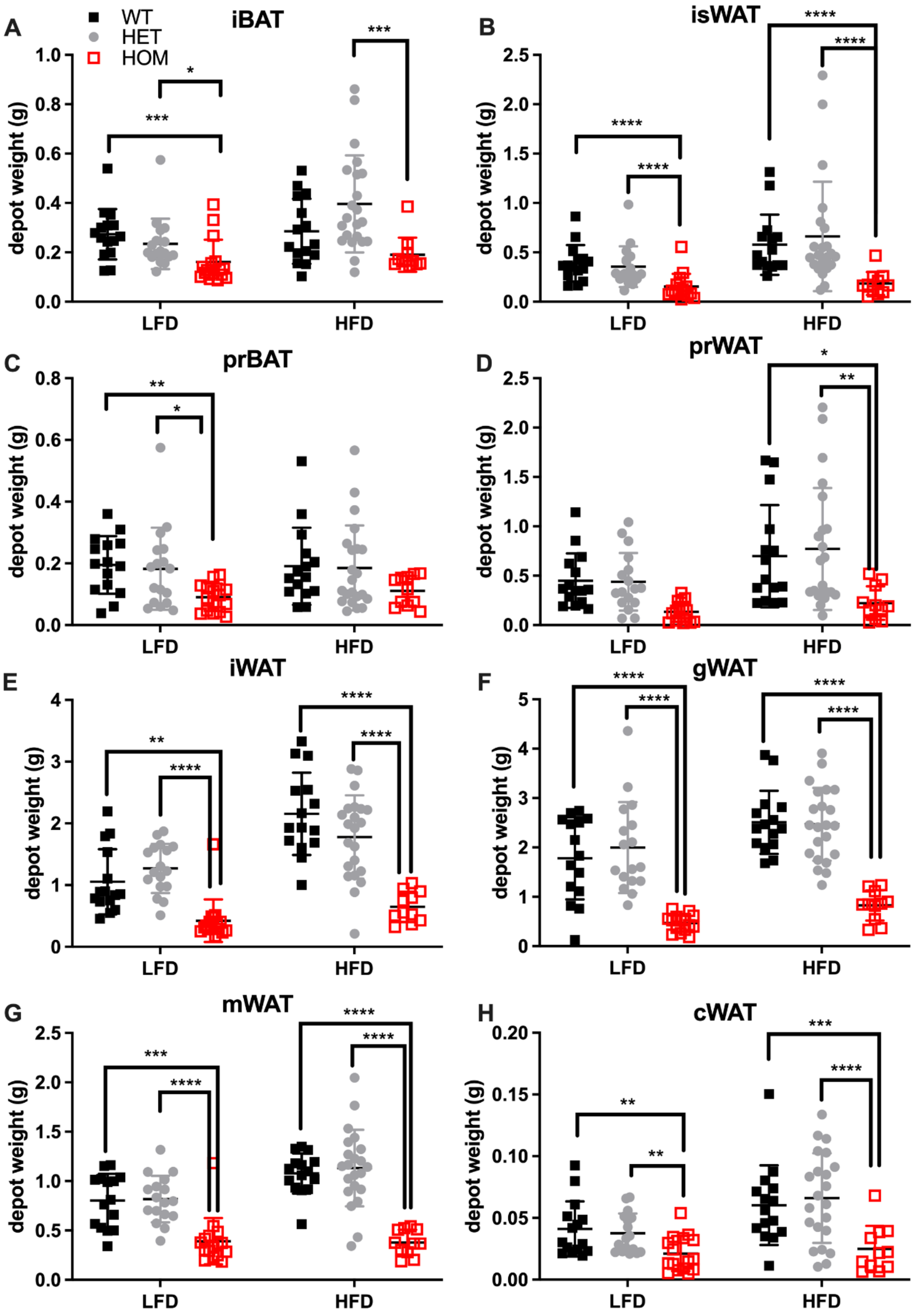

**Supp. Fig. 5 - Multiple female *Wars2*$^{V117L/V117L}$ fat depots are affected.** Fat depots of 6-month old female (n = 11-22) mice on low fat (LFD) or high fat (HFD) diets were dissected and the following fat depots weighed: interscapular BAT (iBAT, A) , interscapular WAT (isWAT, B), perirenal BAT (prBAT, C), perirenal WAT (prWAT, D), inguinal WAT (iWAT, E), gonadal WAT (gWAT, F), mesenteric WAT (mWAT, G), cardiac WAT (cWAT, H). Data in E and F are from Figure 3 and are included here for comparison. One HOM gWAT LFD diet outlier was identified using the Prism ROUT method and excluded. Normality of distribution was evaluated using D'Agostino & Pearson normality test. In order to normalise distribution, data was Y=log2(Y) transformed for iBAT, isWAT, prBAT, isWAT, prWAT and cWAT before analysis. iWAT showed some deviation from normality ($P = 0.0476$) and was analysed as raw data. Significance was tested using 2-way ANOVA with post-hoc Tukey's multiple comparisons tests between all the groups. Significant differences in multiple comparisons of WT, HET and HOM on each diet are depicted as $^*p < 0.05$, $^{**}p < 0.01$, $^{***}p < 0.001$.

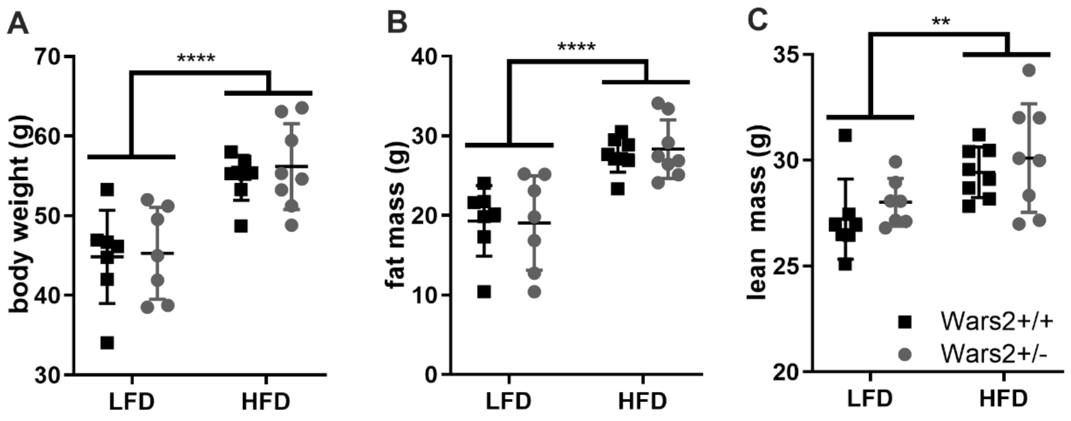

**Supp. Fig. 6 Heterozygous knockout Wars2$^{+/-}$ mice show no difference in bodyweight and body composition.** 12-month old females (n = 7–8) either on low fat (LFD) or high fat (HFD) diets were compared for bodyweight (A), fat mass (B) and lean mass (C). Mean ± SD. Significance was assessed using 2-way ANOVA (diet and genotype) with Sidak's multiple comparisons test.

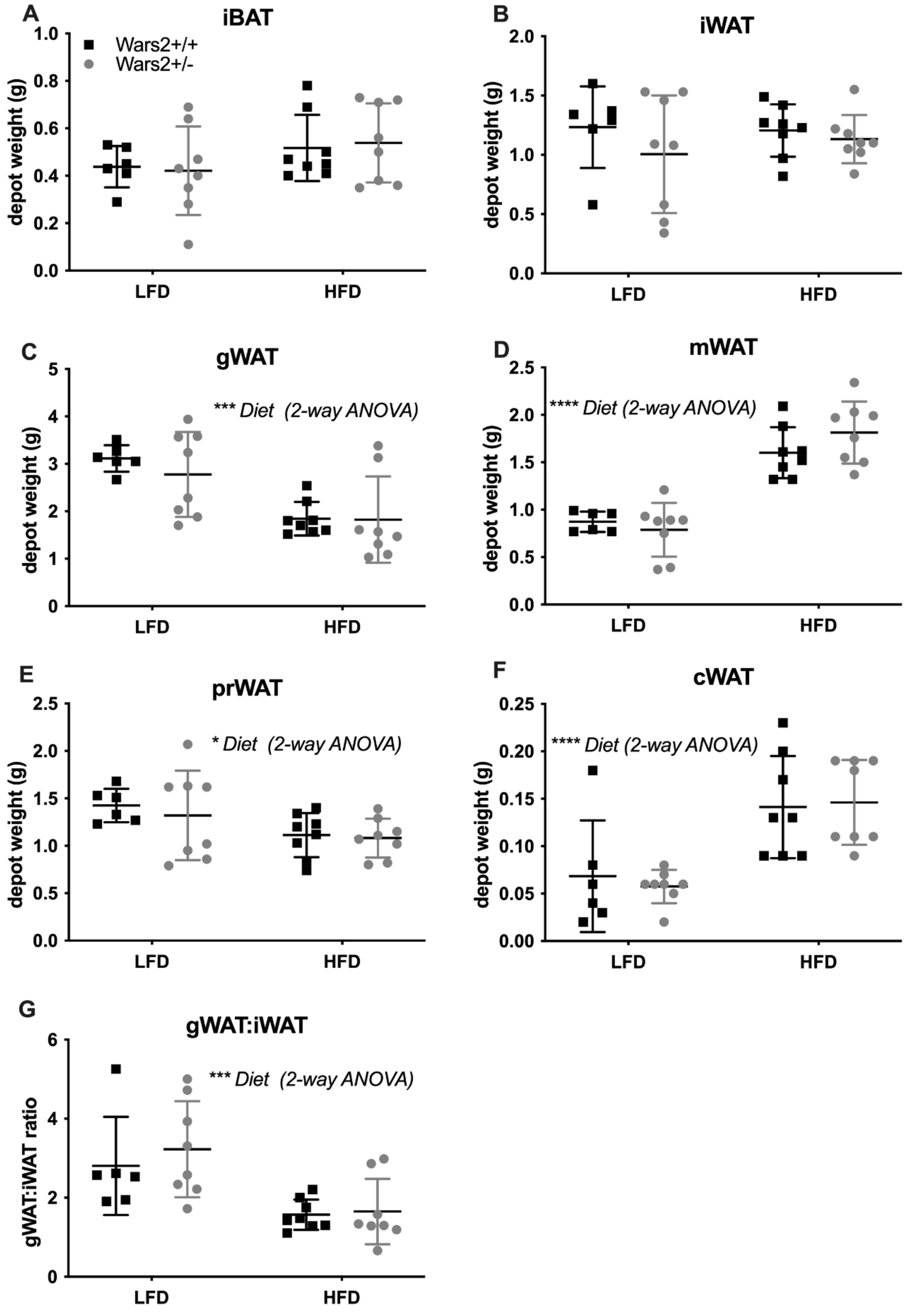

**Supp. Fig. 7 Heterozygous knockout *Wars2*$^{+/-}$ female mice showed no difference in fat depot weights.** 12-month old females (n = 7-9) either on low fat (LFD) and high fat (HFD) diets were sacrificed and the following fat depots were weighed: interscapular BAT (iBAT, A), inguinal WAT (iWAT, B), gonadal WAT (gWAT, C), mesenteric WAT (mWAT, D), perirenal WAT (prWAT, E), cardiac WAT (cWAT, F). Mean ± SD. Normality of distribution was evaluated using D'Agostino & Pearson normality test. In order to normalise distribution, data was Y=log2(Y) transformed for gWAT before analysis. All data was analysed using 2-way ANOVA and significant factors due to diet indicated *$p < 0.05$, **$p < 0.01$, ***$p < 0.001$.

**Supplementary Table 1: A, Factor analysis of area under the curve (AUC) analysis of data in Figure 4 and 5. B, Multiple comparison analysis of area under the curve (AUC) data in Figure 4 and 5.** Area under the curve was calculated on the data presented in Figure 4 and 5, between 6 and 24 weeks of age with the baseline set at zero. Data was not available on all animals at 4 weeks and therefore was not included in the calculation of AUC. For male mice one HOM on a LFD and one WT on a HFD were excluded as outliers (identified using ROUT in GraphPad PRISM 9) and three mice (one HET on LFD, Het on HFD and one WT HFD were excluded due to incomplete data). AUC was analysed using 2way ANOVA to identify the source of variation and multiple comparisons made using Tukey's multiple comparisons test (GraphPad PRISM 9). Lightly shaded boxes are not significant (P = >0.05). LFD, low fat diet; HFD, high fat diet; WT, wildtype (*Wars2*$^{+/+}$); HET, heterozygous (*Wars2*$^{+/V117L}$); HOM, homozygous (*Wars2*$^{V117L/V117L}$).

**A**

| Area under the Curve (AUC) | Male | Female |
|---|---|---|
| **Bodyweight (BW) g** | | |
| Diet (D) | **<0.0001** (HFD v LFD) | **<0.0001** |
| Genotype (G) | **<0.0001**<br>(LFD WT, HET, HOM n = 9,16,16 and HFD WT, HET, HOM n = 14,18,11) | **<0.0001**<br>(LFD WT, HET, HOM n = 15,17,16 and HFD HET, HOM n = 15,22,11) |
| DxG Interaction | **0.0004** | **0.0003** |
| **Fat Mass (FM) g** | | |
| Diet (D) | **<0.0001** | **<0.0001** |
| Genotype (G) | **<0.0001**<br>(LFD WT, HET, HOM n = 9,16,16 and HFD HET, HOM n = 14,19,11) | **<0.0001**<br>(LFD WT, HET, HOM n = 15,17,15 and HFD HET, HOM n = 15,22,12) |
| DxG Interaction | **<0.0001** | **<0.0001** |
| **Lean Mass (LM) g** | | |
| Diet (D) | 0.8740 | 0.9906 |
| Genotype (G) | **<0.0001**<br>(LFD WT, HET, HOM n = 9,16,16 and HFD HET, HOM n = 14,18,11) | **<0.0001**<br>(LFD WT, HET, HOM n = 15,17,16 and HFD HET, HOM n = 15,22,11) |
| DxG Interaction | 0.0571 | 0.1014 |

**B**

| **AUC** | **Male** | | | **Female** | | |
|---|---|---|---|---|---|---|
| | BW | FM | LM | BW | FM | LM |
| **LFD** | | | | | | |
| WT vs. HET | 0.0502 | 0.1198 | 0.0581 | 0.9896 | 0.9998 | 0.9595 |
| WT vs. HOM | **<0.0001** | **<0.0001** | 0.2077 | **0.0031** | **0.0001** | 0.1205 |
| HET vs. HOM | **<0.0001** | **<0.0001** | **<0.0001** | **0.0014** | **<0.0001** | **0.0565** |
| **HFD** | | | | | | |
| WT vs. HET | 0.7843 | 0.9863 | 0.8008 | 0.1492 | 0.2258 | 0.0860 |
| WT vs. HOM | **<0.0001** | **<0.0001** | **<0.0001** | **<0.0001** | **<0.0001** | **0.0111** |
| HET vs. HOM | **<0.0001** | **<0.0001** | **<0.0001** | **<0.0001** | **<0.0001** | **<0.0001** |
